# A Murine Model of Post-acute Neurological Sequelae Following SARS-CoV-2 Variant Infection

**DOI:** 10.1101/2024.01.03.574064

**Authors:** Ankita Singh, Awadalkareem Adam, Aditi, Bi-Hung Peng, Xiaoying Yu, Jing Zou, Vikram V Kulkarni, Peter Kan, Wei Jiang, Pei-Yong Shi, Parimal Samir, Irma Cisneros, Tian Wang

## Abstract

Viral variant is one known risk factor associated with post-acute sequelae of COVID-19 (PASC), yet the pathogenesis is largely unknown. Here, we studied SARS-CoV-2 Delta variant-induced PASC in K18-hACE2 mice. The virus replicated productively, induced robust inflammatory responses in lung and brain tissues, and caused weight loss and mortality during the acute infection. Longitudinal behavior studies in surviving mice up to 4 months post-acute infection revealed persistent abnormalities in neuropsychiatric state and motor behaviors, while reflex and sensory functions recovered over time. Surviving mice showed no detectable viral RNA in the brain and minimal neuroinflammation post-acute infection. Transcriptome analysis revealed persistent activation of immune pathways, including humoral responses, complement, and phagocytosis, and reduced levels of genes associated with ataxia telangiectasia, impaired cognitive function and memory recall, and neuronal dysfunction and degeneration. Furthermore, surviving mice maintained potent T helper 1 prone cellular immune responses and high neutralizing antibodies against Delta and Omicron variants in the periphery for months post-acute infection. Overall, infection in K18-hACE2 mice recapitulates the persistent clinical symptoms reported in long COVID patients and may be useful for future assessment of the efficacy of vaccines and therapeutics against SARS-CoV-2 variants.

## INTRODUCTION

Acute COVID-19 infection typically lasts 4 weeks from symptomatic onset and results in diverse clinical outcome ranging from mild to severe pneumonia, life-threatening multiorgan failure, and death ^1,2^. An estimated 30 to 50% of COVID-19 survivors suffer a post-viral syndrome known as long-COVID [post-acute sequelae of COVID-19, (PASC)], which encompasses ongoing persistent symptoms and complications beyond the initial 4 weeks of infection ^3,4^. The key features of long-COVID include neurological symptoms, such as fatigue, cognitive dysfunction (brain fog, memory loss, attention disorder), and sleep disturbances ^5,6^. Psychiatric manifestations (anxiety and depression) are also common ^7,8^. Although the severity of acute infection is considered as one major risk factor for developing PASC ^9,10^, increasing evidence suggests that long-COVID also occurs in people with non-symptomatic or non-hospitalized status during acute infection ^11,12^. PASC has posed a significant threat to the global healthcare system. Nevertheless, our current understanding of its underlying mechanisms is limited.

Acute SARS-CoV-2 infection has been studied in various animal models, including hamsters, ferrets, non-human primate (NHP)s, rats, and mice ^13,14^. Mice are easier to work with, and most amenable to immunological manipulation. Angiotensin converting enzyme 2 (ACE2) is the cell entry receptor for SARS-CoV-2 ^15^, and mouse ACE2 shows key differences from human ACE2 (hACE2), thus wild-type immunocompetent mice present challenges for infection with human SARS-CoV2 variants. To overcome this, delivery of adeno-associated virus (AAV)–mediated expression of hACE2 into the respiratory tract or use of the mouse-adapted SARS-CoV-2 (SARS-CoV-2 MA strain or CMA strain), which incorporates key mutations that allows the virus to utilize mouse ACE2 for entry into cells in immunocompetent mice, results in a productive infection with mild acute respiratory distress syndrome ^16–18^. Acute SARS-CoV-2 infection in K18-hACE2 transgenic mice, which express hACE2 in lung epithelial cells also induces weight loss, interstitial pneumonitis, encephalitis, and death^19–22^. In this study, we infected K18-hACE2 mice with SARS-CoV-2 Delta variant and performed longitudinal behavior analysis for up to 4 months post-acute infection. Mice surviving acute Delta variant infection displayed persistent abnormalities in neuropsychiatric state and motor behavior post-acute infection. Although surviving mice developed and maintained potent Th1-prone cellular and antibody responses in the periphery, transcriptome analysis suggested persistent activation of immune pathways in the CNS, and cognitive and neuronal dysfunction for months post-acute infection.

## Materials and Methods

### Viruses

SARS-CoV-2 Delta and Omicron B.A.2 strains were obtained from the World Reference Center for Emerging Viruses and Arboviruses (WRCEVA) at the University of Texas Medical Branch (UTMB) and were amplified twice in Vero E6 cells.

### SARS-CoV-2 infection in mice

6-to 8-week-old K18-hACE2 mice (Jackson Lab, stock #034860) were bred and maintained at UTMB animal facility. All animal experiments were approved by the Animal Care and Use Committees at UTMB. Mice were infected intranasally (i.n) with 600 plaque and 800 forming unit (PFU) of SARS-CoV-2 Delta or Omicron B.A.2 strain respectively. Infected mice were monitored twice daily for morbidity and mortality. In some experiments, on days 3, 6, 7, 8, 9, 1-month, 2-months, and 4-months post infection (pi), mice were euthanized for tissue collection for viral load, histopathology analysis, and immune cell function studies.

### Quantitative PCR (Q-PCR)

The sequences of the primer sets and PCR reaction conditions were described previously ^23–25^ and in **Table S1**. Tissues were resuspended in TRIzol for RNA extraction according to the manufacturer’s instructions (Thermo Fisher). RNA concentration and purity was determined by using WPA Biowave DNA Spectrophotometer. Complementary (c) DNA was then synthesized by using a qScript cDNA synthesis kit (Bio-Rad). Expression of SARS-CoV-2 S gene and mouse inflammatory cytokine and chemokine genes (IL-1β, IL-6, TNF-α, CCL2, CCL5, CCL7, CXCL10, and CCL11) were measured by Q-PCR using CFX96 real-time PCR system (Bio-Rad). PCR cycling conditions were as follows: 95°C for 3 min, 45 cycles of 95°C for 15 s, and 60°C for 1 min. Gene expression was calculated using the formula 2 -[Ct(target gene)-Ct(β-actin)] as described before ^26^.

### RNAseq and gene set enrichment analysis and Cytoscape analysis

RNA was extracted from brain tissues as described above and 1-3 µg of RNA of each sample was used for RNAseq analysis. The RNA quality was verified by the Next Generation Sequence (NGS) core laboratory using a Nanodrop ND-1000 spectrophotometer (Thermofisher) and an Agilent Bioanalyzer 2100 (Agilent Technologies, Santa Clara, CA). PolyA+ RNA was purified from ∼100 ng of total RNA and sequencing libraries were prepared with the NEBNext Ultra II RNA library kit (New England Biolabs) following the manufacturer’s protocol. Libraries were pooled and sequenced on an Illumina NextSeq 550 High Output flow-cell with a paired-end 75 base protocol. Pathway enrichment analysis was performed using GSEA version 3.0 ^27,28^. Specifically, analysis parameters were set to 2000 gene set permutation and gene set size limit 15-500. Primary gene sets investigated were obtained from the David Bader lab (http://download.baderlab.org/EM_Genesets/). GSEA FDR Q < 0.05 cutoff was applied to examine enriched gene sets in our dataset. Cytoscape Enrichment map tool was used to visualize results from the GSEA analysis. GSEA output files were uploaded in the Enrichment Map app of Cytoscape and FDR Q value cut off was set to 0.01. AutoAnnotate Cytoscape app was used to define the clusters automatically.

### Plaque assay

Vero E6 cells were seeded in 6-well plates and incubated at 37°C. Tissue homogenates were serially diluted (10-fold) in DMEM with 2% FBS and 0.2 ml was used to infect cells at 37°C for 1 h. After incubation, samples were overlaid with MEM (Gibco) with 8% FBS and 1.6% agarose (Promega). After 48 h, plates were stained with 0.05% neutral red (Sigma-Aldrich) and plaques were counted to calculate virus titers expressed as PFU/ml.

### Histopathology studies

Lung and brain tissues were fixed in formalin (Thermo scientific) for at least two days before embedment in optimal cutting temperature compound. Hematoxylin and eosin (H&E) staining was performed at the Histopathology Laboratory Core of UTMB.

### Behavioral studies

At 1, 2, 3, and 4 months pi, all surviving animals underwent neurological assessments using a modified SmithKline Beecham, Harwell, Imperial College, Royal London Hospital phenotype assessment (SHIRPA) protocol. Mice were also weighed at the above time points to confirm their growth. For the modified SHIRPA assessment, each mouse was placed in a transparent cylindrical viewing jar and observed for 5 min. Observations in body position, spontaneous activity, respiration rate, tremor, and defecation were noted. Subsequently, the mouse was transferred to an open field arena at which time transfer arousal and gait were noted. Following transfer, the mice were allowed to freely move around the open field arena for 30 sec and the number of times that all four limbs crossed into new quadrants was counted to evaluate locomotor activity. For the next 5 min, gait, eye opening, piloerection, pelvic elevation, tail position, and touch escape were observed in the open field arena. Further, tail lifting was performed to evaluate trunk curl and visual placing followed by assessment of reach touch, grip strength, whisker response, palpebral reflex, and ear twitch above arena. Additional behaviors of each mouse, such as fear, biting, irritability, and aggression, were observed throughout the procedure. Based on the observation, scores were provided (0, 1, 2, or 3), 2 was considered normal behavior, any score outside of 2 was considered abnormal behavior. Each parameter assessed by SHIRPA were grouped into five functional categories (see **Table 1**). To measure motor coordination, a parallel rod floor test was performed. Briefly, mice were placed in the center of the cage that was covered with horizontal rods for 2 min. Foot/paw slips were counted and recorded manually.

**Table 1:**
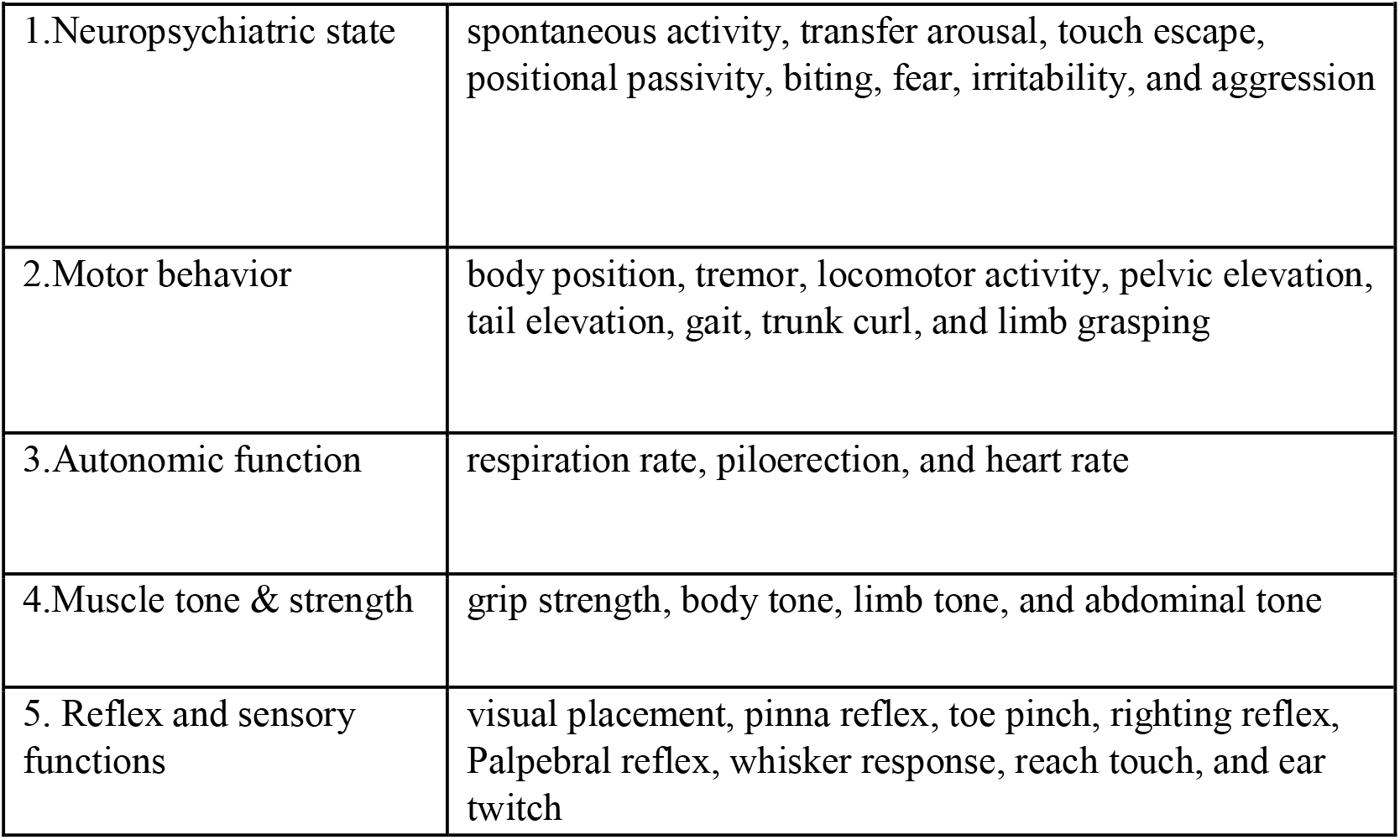
Modified SHIRPA Assessment.

### Immunofluorescence staining

Formalin fixed, paraffin-embedded tissue blocks were cut into 10 μm sections, which were deparaffinized with Xylene and, rehydrated with ethanol/water. To detect Iba1 and GFAP expression, tissue sections were exposed to anti-Iba1 rabbit antibody (FUJIFILM Wacko, 013-27691, 1:300 dilution) and anti-GFAP mouse antibody (Sigma, G3893, 1:300 dilution) for overnight at room temperature, respectively. After washing with phosphate-buffered saline (PBS), slides were exposed to goat anti-rabbit Alexa flour Plus 555 (ThermoFisher, A32732, 1:1000 dilution) and goat anti-mouse Alexa flour 488 (ThermoFisher, A32723, 1:1000 dilution) secondary antibodies for 1 h at room temperature. For nuclei staining, we used 4′,6′-diamidino-2-phenylindole DAPI (D9542, Sigma). Slides were washed with PBS and mounted with ProLong Gold Antifade (ThermoFischer, P36930). Images were captured using Olympus BX53 microscope.

### Antibody and SARS-CoV-2 spike (S) protein ELISA

ELISA plates were coated with 100 ng/well recombinant SARS-CoV-2 RBD protein (RayBiotech) for overnight at 4°C. The plates were washed twice with PBS containing 0.05% Tween-20 (PBS-T) and then blocked with 8% FBS for 1.5 h. Sera were diluted 1:100 in blocking buffer and were added for 1 h at 37°C. Plates were washed 5 times with PBS-T. Goat anti-mouse IgG (Sigma) coupled to horseradish peroxidase (HRP) or alkaline phosphatase was added at a 1:2000 dilutions for 1 h at 37°C. This was followed by adding TMB (3, 3, 5, 5′-tetramethylbenzidine) peroxidase substrate (Thermo Fisher Scientific) for about 10 min. The reactions were stopped with 1M sulfuric acid, and the intensity was read at an absorbance of 450 nm.

For SARS-CoV-2 S protein ELISA, plates were coated with 50 ng/well (5 µg/ml in coating buffer) of diluted capture antibody (Thermo Fisher Scientific, USA) overnight at 4°C. The plates were washed twice with PBS-Tween (PBS-T) and then blocked with 8% FBS for 1.5 h. Standard (recombinant SARS-CoV-2 S protein, Sino Biological, USA) was diluted in blocking buffer and the brain suspension were added for 1 h at 37°C. Plates were washed 5 times with PBS-T. SARS-CoV-2 S1 protein primary antibody (Thermo Fisher Scientific) at 1µg/ml were added for 1 h at 37°C. Plates were again washed 5 times with PBS-T. Goat anti-Rabbit IgG (Thermo Fisher Scientific) coupled to horseradish peroxidase (HRP) was added at a 1:2000 dilutions for 1 h at 37 °C, followed by adding TMB (3, 3, 5, 5′-tetramethylbenzidine) peroxidase substrate (Thermo Fisher Scientific) for 30 min. The reactions were stopped by 1M sulfuric acid, and the intensity was read at an absorbance of 450 nm.

#### IFN-γ ELISPOT

Millipore ELISPOT plates (Millipore Ltd) were coated with anti-mouse IFN-γ capture Ab at 1:100 dilution (Cellular Technology Ltd) at 4°C overnight. Splenocytes were stimulated with SARS-CoV-2 S peptide pools (2 μg/ml, Miltenyi Biotec) for 24 h at 37°C. Cells were stimulated with anti-CD3 (1 μg/ml, e-Biosciences) or medium alone, as controls. This was followed by incubation with biotin-conjugated anti-IFN-γ at 1:100 dilution (Cellular Technology Ltd,) for 2 h at room temperature, followed by incubation with alkaline phosphatase-conjugated streptavidin for 30 min. The plates were washed and scanned using an ImmunoSpot 6.0 analyzer and analyzed by ImmunoSpot software to determine the spot-forming cells (SFC) per 10^6^ splenocytes.

### Intracellular cytokine staining (ICS)

Splenocytes were incubated with SARS-CoV-2 S peptide pools (1μg/ml, Miltenyi Biotec) for 6 h in the presence of GolgiPlug (BD Bioscience). Cells were harvested and stained with antibodies for CD3, CD4, or CD8, fixed in 2% paraformaldehyde, and permeabilized with 0.5% saponin before adding anti-IFN-γ (Thermo Fisher). Samples were acquired by a C6 Flow Cytometer instrument. Dead cells were excluded based on forward and side light scatter. Data were analyzed with a CFlow Plus Flow Cytometer (BD Biosciences).

### Fluorescent focus reduction neutralization test

Neutralization titers of mice sera were measured by a fluorescent focus reduction neutralization test (FFRNT) using the mNG reporter SARS-CoV-2 as previously reported with some modifications ^29^. Briefly, Vero E6 cells (2.5 × 10^4^) were seeded in each well of black μCLEAR flat-bottom 96-well plate (Greiner Bio-one™). The cells were incubated overnight at 37°C with 5% CO_2_. On the following day, each serum was 2-fold serially diluted in the culture medium with the first dilution of 1:20. Each serum was tested in duplicates. The diluted serum was incubated with 100-150 fluorescent focus units (FFU) of mNG Delta and BA.2 SARS-CoV-2 at 37°C for 1 h (final dilution range of 1:20 to 1:20480), respectively. After that, the serum-virus mixtures were inoculated onto the pre-seeded Vero E6 cell monolayer in 96-well plates. After 1 h of infection, the inoculum was removed and 100 μl of overlay medium (DMEM supplemented with 0.8% methylcellulose, 2% FBS, and 1% P/S) was added to each well. After incubating the plates at 37°C for 16 h, raw images of mNG fluorescent foci were acquired using CytationTM 7 (BioTek) armed with 2.5× FL Zeiss objective with wide field of view and processed using the software settings (GFP [469,525] threshold 4000, object selection size 50–1000 µm). The foci in each well were counted and normalized to the non-serum-treated controls (set as 100%) to calculate the relative infectivity. The neutralizing titer 50 (NT_50_) was calculated manually as the highest dilution of the serum sample that prevents at least 50% fluorescence foci formation in infected cells. A titer is calculated for each of the two replicates of a sample and the geometric mean titer (GMT) of the two is reported as the final sample titer.

### Statistical analysis

Survival curve comparison was performed using GraphPad Prism software 9.4.1, which uses the log-rank test. Values for viral load, cytokine production, antibody titers, and T cell response experiments were compared using Prism software statistical analysis and were presented as means ± SEM. *P* values of these experiments were calculated with a non-paired Student’s t test. Parameters of behavior changes at month 1 and results of parallel rod test were compared using Student’s t test. For the categorical, longitudinal measures of each parameter in modified SHIRPA testing, we considered a score of 2 as normal activity and the other scores (0, 1, and 3) as abnormal activity. For selected parameters, changes were presented over time in a stacked bar chart with Sankey-style overlays using SAS version 9.4 (SAS Inc., Cary, NC). All tests were two-sided with a significance level of 0.05.

## RESULTS

### SARS-CoV-2 variant replicated in brain and lung tissues and induced inflammatory responses in both tissues during the acute infection in K18-hACE2 mice

Several reports suggest that the frequencies of PASC symptoms increased with SARS-CoV-2 variants, in particular, the pre-Omicron variant compared to the original prototype virus infection ^3,30–32^. Thus, to investigate the mechanisms of PASC, we infected K18-hACE2 mice with SARS-CoV-2 Delta variant. Viral load analysis, survival/weight monitoring, immunological and histopathology studies, and behavior assessment were performed at both acute infection and at 1 to 4 months post-acute COVID infection (**Fig. 1A**). Initially, 6-to 8-week-old K18-hACE2 mice were intranasally (i.n.) inoculated with a sublethal dose of SARS-CoV-2 Delta variant strain and monitored daily for morbidity and mortality. Infected mice exhibited weight loss starting on day 6 pi and succumbed to infection as early as day 7. About 22% of mice infected with the Delta variant survived the 4-week pi interval (**Fig. 1B-C**). In the brain, viral RNA but not infectious virus was detected at day 3. Viral loads increased significantly at day 6. Viral RNA levels decreased after day 6 but were continuously detectable at days 7, 8, and 9 pi (**Fig. 1D-E**, **Supplementary Fig. 1A**). At days 3 and 6, proinflammatory cytokines, including IL-1β, IL-6, and TNF-α (**Fig. 1F**), and chemokines, such as CCL2, CCL5, CXCL10, and CCL11, were induced in the brains of infected mice (**Fig. 1G**). Histopathology analysis also revealed viral encephalitis with perivascular infiltrations and microglial activation in the cortex of infected mice, but not in mock-infected mice, which together suggest neuroinflammation induction in Delta variant-infected mice (**Fig. 1H**). In the lung, viral loads were high at day 3 but diminished at day 6 (**Fig. 2A-B**). Viral RNA remained detectable in the lungs at days 7, 8, and 9 pi (**Supplementary Fig. 1B**). Proinflammatory cytokines and chemokines, including IL-1β, IL-6, CCL2, CCL5, CXCL10, and CCL11, were triggered in the lungs of infected mice compared to the mock (**Fig. 2C-D)**. Lung pathology study showed mononuclear cell infiltration in peribronchiolar and perivascular areas as well as in the alveolar septa of the infected mice, but not in the mock group (**Fig. 2E**). Overall, these results suggest that SARS-CoV-2 Delta variant replicated in lung and brain tissues and triggered inflammation during the acute infection phase in K18-hACE2 mice.

**Figure 1.**
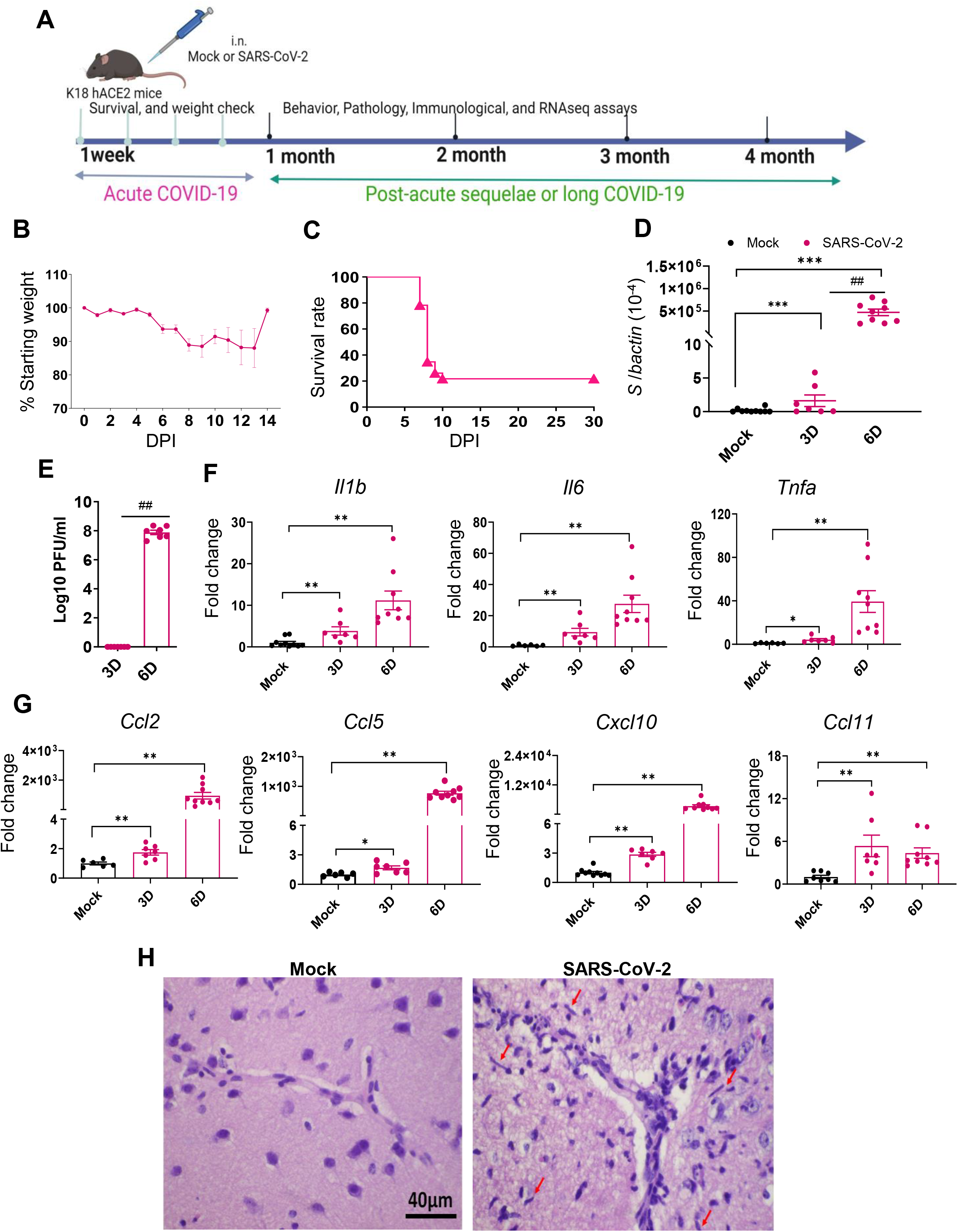
SARS-CoV-2 Delta variant replicated and induced inflammatory responses in brain tissues during acute infection in K18-hACE2 mice. 6-to 8-week-old K18-hACE2 mice were i.n. infected with a sublethal dose of SARS-CoV-2 Delta variant strain (or mock infected) and monitored daily for morbidity and mortality. A. Study design. B. Mouse weight loss. Weight loss is indicated by percentage using the weight on the day of infection as 100%. n= 23. C. Survival rate. D-E. SARS-CoV-2 viral loads in brain were measured by Q-PCR (D) and plaque assay (E) at indicated days (D) pi. F-G. Cytokine (F) and chemokines (G) expression levels in the brains were measured by Q-PCR. Data are presented as the fold increase compared to mock-infected mice (means ± SEM). n= 7 to 10. H. Histopathology of brains of Delta variant -infected mice revealed viral encephalitis with perivascular infiltrations and microglial activation (arrows) in the cortex, but not in mock-infected mice \*\*\**P* < 0.001, \*\**P* < 0.01, or \**P* < 0.05 compared to mock. ^##^*P* < 0.01 compared to 3D.

**Figure 2.**
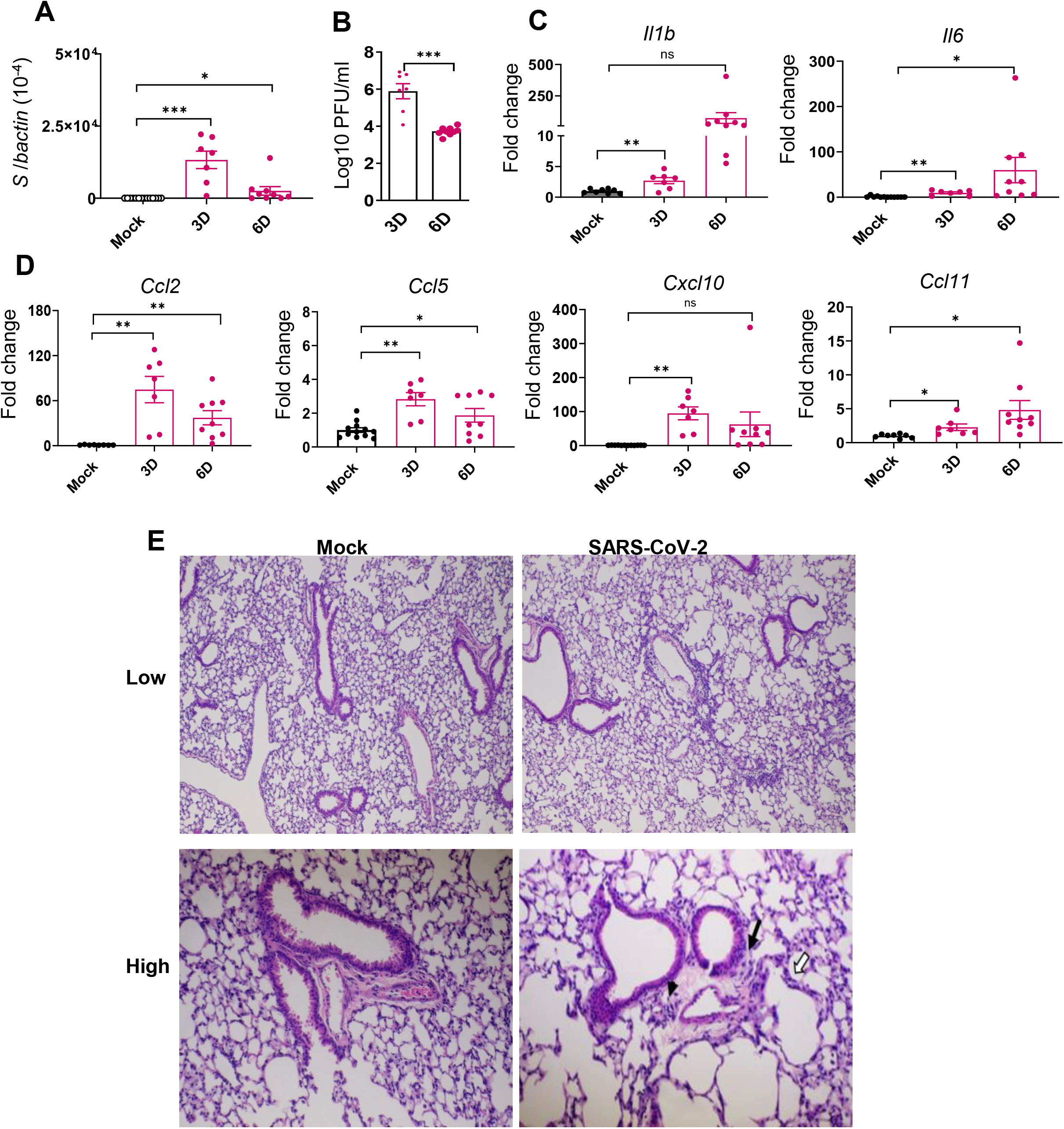
SARS-CoV-2 Delta variant replicated in the lung tissues and induced inflammatory responses in K18-hACE2 mice during acute infection. 6-to 8-week-old K18-hACE2 mice were i.n. infected with a sublethal dose of SARS-CoV-2 Delta variant strain or mock infected. Viral loads in lung tissues were measured by Q-PCR (**A**) and plaque assays (**B**) at indicated days (D) pi. **C-D**. Cytokine (**C**) and chemokines (**D**) expression levels in the lung were measured by Q-PCR. Data are presented as the fold increase compared to mock-infected mice (means ± SEM). n= 7 to 9. **E.** Histopathology of Delta variant-infected (right panels) or mock (left panels) -infected lungs. Low power views of representative areas (top panels) from each group show moderate inflammation in the infected mice. At higher power views (low panels), mononuclear cell infiltrations are observed in peribronchiolar (black arrow) and perivascular (arrowhead) areas as well as in the alveolar septa (white arrows). Bar = 200μm for top panels; Bar = 80 μm for low panels. \*\*\**P* < 0.001, \*\**P* < 0.01, or \**P* < 0.05 compared to mock.

### SARS-CoV-2 Delta variant-infected mice displayed neurological behavior changes months post-acute infection

Based on National Institute for Health and Care Excellence guidelines, PASC is displayed at 4 weeks or more after the start of acute COVID-19 infection ^3^. Here, we did not detect infectious virus (data not shown) nor viral RNAs (**Supplementary Fig. 1A-B**) at 1 and 4 -month pi in the brain and lung tissues. No significant levels of viral S1 protein were detected in the brain tissues beyond 1 month, which together indicate viral clearance at the post-acute phase (**Supplementary Fig. 1C**). Viral loads in several other periphery tissues, including liver, kidney, and blood, were also measured during acute and post-acute infection. No detectable viral RNA was found in these tissues except at day 6 in the kidneys (**Supplementary Fig. 1D-F**). Histopathological analysis revealed no changes in the various regions of brain in the Delta variant-infected mice at 1 or 4-month pi compared to the mock group (**Supplementary Fig. 2A**). In the lung, there were notably increased levels of CCL7 and CXCL-10 at 1 month, and increased levels of IL-6, CCL5, CXCL10, and CCL11 at 4 months in the infected mice compared to the mock group (**Supplementary Fig. 2B)**, though no changes in the levels of IL-1β, TNF-α, and CCL2 were observed (**data not shown**). Thus, viral infection was cleared in the periphery and CNS tissues post-acute infection, and this was accompanied by mild and minimal local inflammatory responses in the lung and brain tissues, respectively.

To assess the infection impact on animals, behavior tests were performed in the surviving mice at 1 month pi using a modified Smith-Kline Beecham, Harwell, Imperial College, Royal London Hospital, phenotype assessment (SHIRPA) protocol ^33,34^. Mock-infected mice were used as controls. The assay involves a battery of semi-quantitative tests for general health and sensory function, baseline behaviors, and neurological reflexes. The individual parameters assessed by SHIRPA were grouped into five functional categories (**Table 1**): 1) Motor behavior test includes body position, tremor, locomotor activity, pelvic elevation, tail elevation, gait, trunk curl, and limb grasping; 2) Autonomic function test includes respiration rate, palpebral closure, and piloerection; 3) Muscle tone and strength includes grip strength, body tone, limb tone; 4) Neuropsychiatric state includes spontaneous activity, transfer arousal, touch escape, positional passivity, biting, fear, irritability, and aggression; 5) Reflex and sensory functions include parameters such as visual placement, toe pinch, and righting reflex ^35^. The SHIRPA assay results showed that mice surviving acute infection with SARS-CoV-2 Delta variant displayed abnormalities mainly in neuropsychiatric state, motor behavior, autonomic function, and reflex and sensory function, compared to the mock group **(Fig. 3A**). Weight loss was not noted in the surviving mice at 1-month pi nor was detected during the rest of 4-month pi interval (**Fig. 3B**). Furthermore, the Delta variant-infected mice showed higher number of foot slips compared to the mock group at 3 month pi in a parallel rod test, which indicates ataxia in the infected mice ^36^ (**Fig. 3C**).

**Figure 3.**
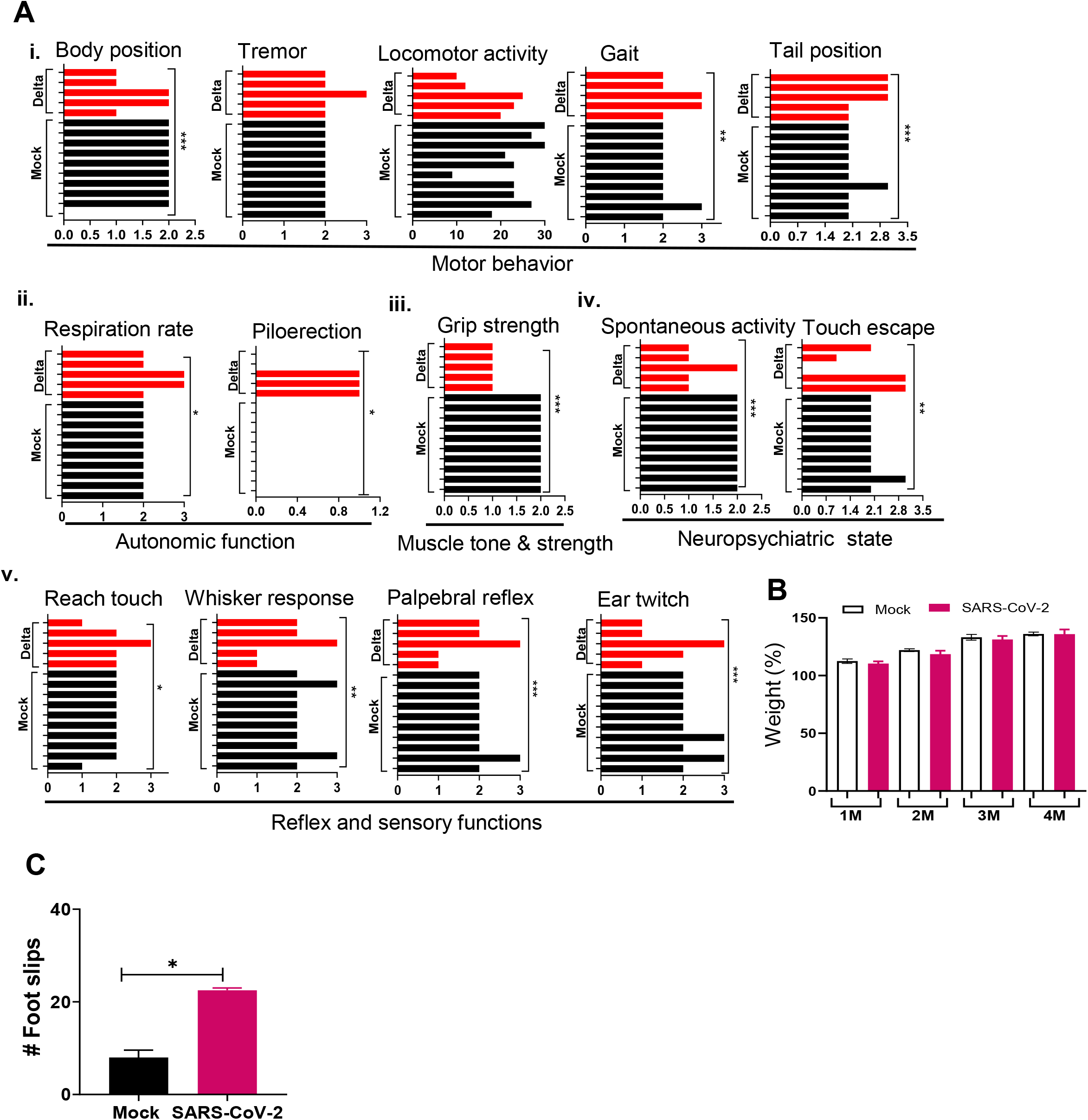
SARS-CoV-2 Delta variant induces behaviors changes post-acute infection. K18-hACE2 mice were infected with a sublethal dose of SARS-CoV-2 Delta variant, or PBS (mock). At 1 month pi or after, surviving mice (n= 10 for mock and n= 5 for SARS-CoV-2 Delta variant infected mice) were assessed for behavior changes via SHIRPA (**A**), weight changes (**B**), and a parallel rod floor test (**C-D**). **A.** At 1 month pi, SARS-CoV-2 variant infected mice showed impaired performance in the SHIRPA assessment. i. Motor behavior; ii. Autonomic function; iii. Muscle tone & strength; iv. Neuropsychiatric state; v. Reflex and sensory functions. **B.** Weight changes during the 4 month pi interval as presented as percentage using the weight on the day of infection as 100%. **C**. Parallel rod floor test. At 3 months pi, surviving mice were placed in the center of the cage coated with horizontal rods for 2 min. Foot/paw slips were counted. \*\*\**P* < 0.001, \*\**P* < 0.01, or \**P* < 0.05 compared to the mock group.

To further assess the impact of SARS-CoV-2 infection on behavioral changes of surviving mice, SHIRPA analysis was performed longitudinally over the 4-month pi period. It was noted that in mice surviving Delta variant infection, the abnormal rates for the parameters such as body position, grip strength, touch escape, and reach touch remained unchanged during the 4-month period (**Fig. 4A**). However, abnormal levels for gait, whisker response, ear twitch, and palpebral reflex decreased over time (**Fig. 4B**). In contrast, abnormalities for parameters including spontaneous activity, tail position, and tremor increased over the 4-month period (**Fig. 4C**). Overall, the neuropsychiatric state and motor behavior of Delta-variant infected mice remained impaired or even deteriorated over months pi; whereas reflex and sensory functions appeared to recover over time.

**Figure 4.**
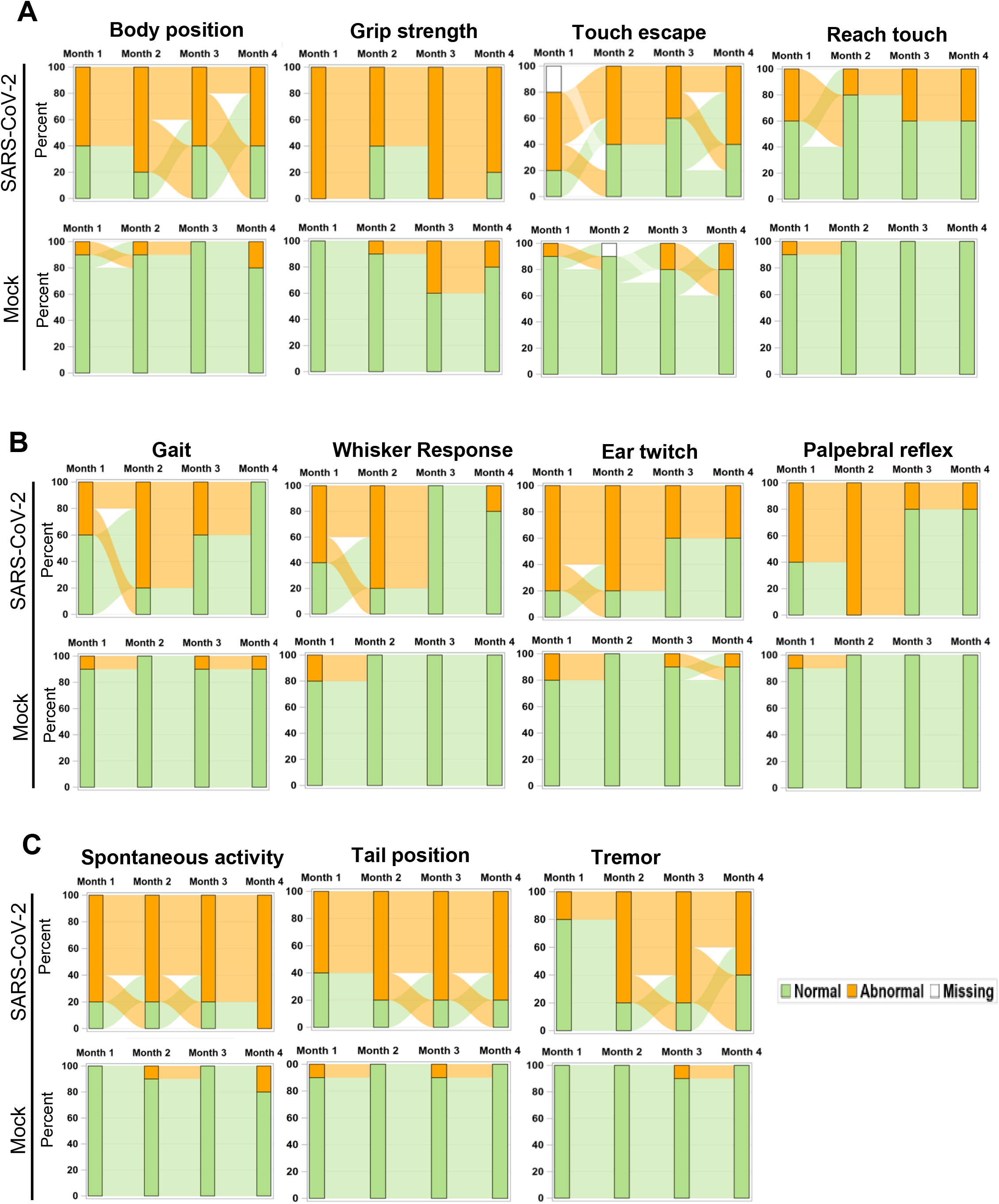
Longitudinal analysis of behavior changes 4 months post-acute infection in mice. Sankey Bar Charts were used to present results of SHIPA analysis. Functional status was defined as abnormal vs. normal and used to form the groups in the stacked bars at each time. Longitudinal stacked bar chart with Sankey-style overlays visualizes how the mice transit between abnormal and normal status over time. The y-axis is the percent in each group and x-axis is the time for the measures collected. The link (bend) between the bars shows the transitions between two states over time. **A.** Parameter abnormality patterns not changed within 4-month pi period. **B-C**. Parameter abnormality rates decreased (**B**) or increased (**C**) within the 4 month pi period.

### Transcriptome analysis revealed persistent activation of immune pathways in the CNS and cognitive and neuronal dysfunction months post-acute infection, though minimal microglia activation was observed. Potent Th1-prone cellular and antibody responses persisted in the periphery post-acute infection

To identify the host factors contributing to SARS-CoV-2 variant-induced PASC, we determined gene expression alterations by comparing the transcriptomes of mock and SARS-CoV-2 Delta variant-infected mouse brains using bulk RNA-seq. Transcriptome analysis identified 3,481 and 18 differentially expressed genes (FDR <0.1) in 1-month and 4-month post-SARS-CoV-2 variant-infected mouse brains, respectively, compared to mock. We then performed gene set enrichment analysis (GSEA) to identify the biological processes that play a role in host response against SARS-CoV-2 infection. GSEA analysis of RNAseq data identified pathways related to immune signaling, such as the “complement activation pathway” and “phagocytosis recognition” (FDR <0.01) as the top enriched pathways (**Fig. 5A**, **Supplementary Fig. 3A**). Interestingly, we also observed enrichment of the same immune signaling related pathways “complement activation pathway,” “phagocytosis recognition,” and “humoral immune response mediated by circulating immunoglobulin” (FDR <0.01) among the top 5 enriched pathways at 4-month pi (**Fig. 5B, Supplementary Fig. 3B**). Notably, the normalized enrichment scores (NES) were higher for these pathways compared to the data set at 1-month pi, an indication of persistence of immune activation in the CNS caused by infiltering immune cells and factors following the initial viral infection. Next, we utilized Cytoscape Enrichment Map and AutoAnnotate tools to identify biological networks that are associated with the enriched gene sets and found immune response cluster in both 1-month and 4-month pi data sets. In addition, other enriched gene sets were annotated as “SARS-CoV-2 translation,” “electron transport process,” and “ribosomal small subunit” indicating that overall nervous system homeostasis is perturbed (**Fig. 5C-D**). Immunofluorescence staining was next performed to detect activation of microglia and astrocytes at both acute infection (day 6) and post-acute infection (1-month and 4-month pi). Microglia activation with increased cell processes was noted at day 6; however, minimal to mild activation of these cells were observed at 1-month and 4-month pi (**Fig. 6A**). Minimal astrocyte activation was detected at 1 month and 4 months pi (data not shown). These data suggest that infiltrating immune cells, not the residential immune cells contribute to neuronal dysfunction at the post-acute stage. Q-PCR analysis was next utilized to determine 8 differentially expressed genes identified by transcriptomic analysis. Reduced levels of expression of *Ddit4*, *Slc38a*2, *Tmem267m*, *Lrrc8c*, and *setd7* genes were noted at 4-month pi, which are associated with ataxia telangiectasia neurodegenerative disease, impairment of memory, synaptic plasticity, motor, and cognitive abilities, neuronal dysfunction and degeneration, and cerebral ischemic stroke respectively^37–40^ (**Fig. 6B**).

**Figure 5.**
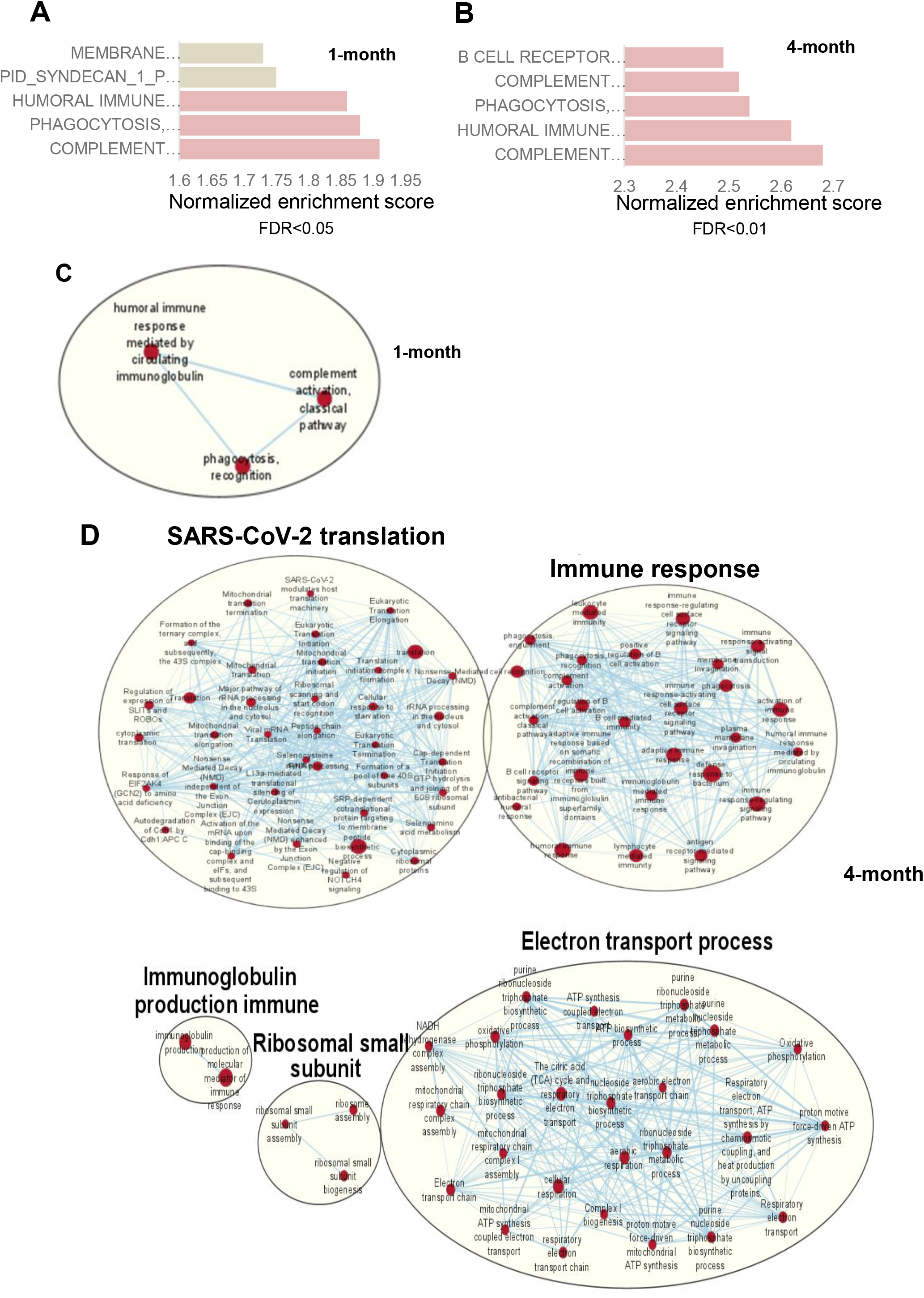
RNAseq analysis of SARS-CoV-2-infected mouse brain shows genes with upregulation of immune signaling and enrichment of immune signaling related pathways. **A.** Pathway enrichment analysis of differential expressed genes using GSEA shows the top 5 upregulated pathways in SARS-CoV2 infected mouse brain at 1 month (**A**) and 4 months (**B**) compared to mock treated brain. Gray bar represents FDR >0.05 while orange bars represent FDR <0.05 **C-D.** Cytoscape enrichment map (FDR *Q* value < 0.01) of GSEA pathways enriched in upregulated genes in SARS-CoV-2 infected mouse brain tissue at 1 month (**C**) and 4 months (**D**) pi compared to mock-treated sample. Clusters of nodes were labeled using Auto Annotate feature of Cytoscape application. Red nodes represent upregulated gene set enrichment and their node size represents the gene-set size. The thickness of line connecting the nodes represents the degree of overlap between two gene-sets.

**Figure 6.**
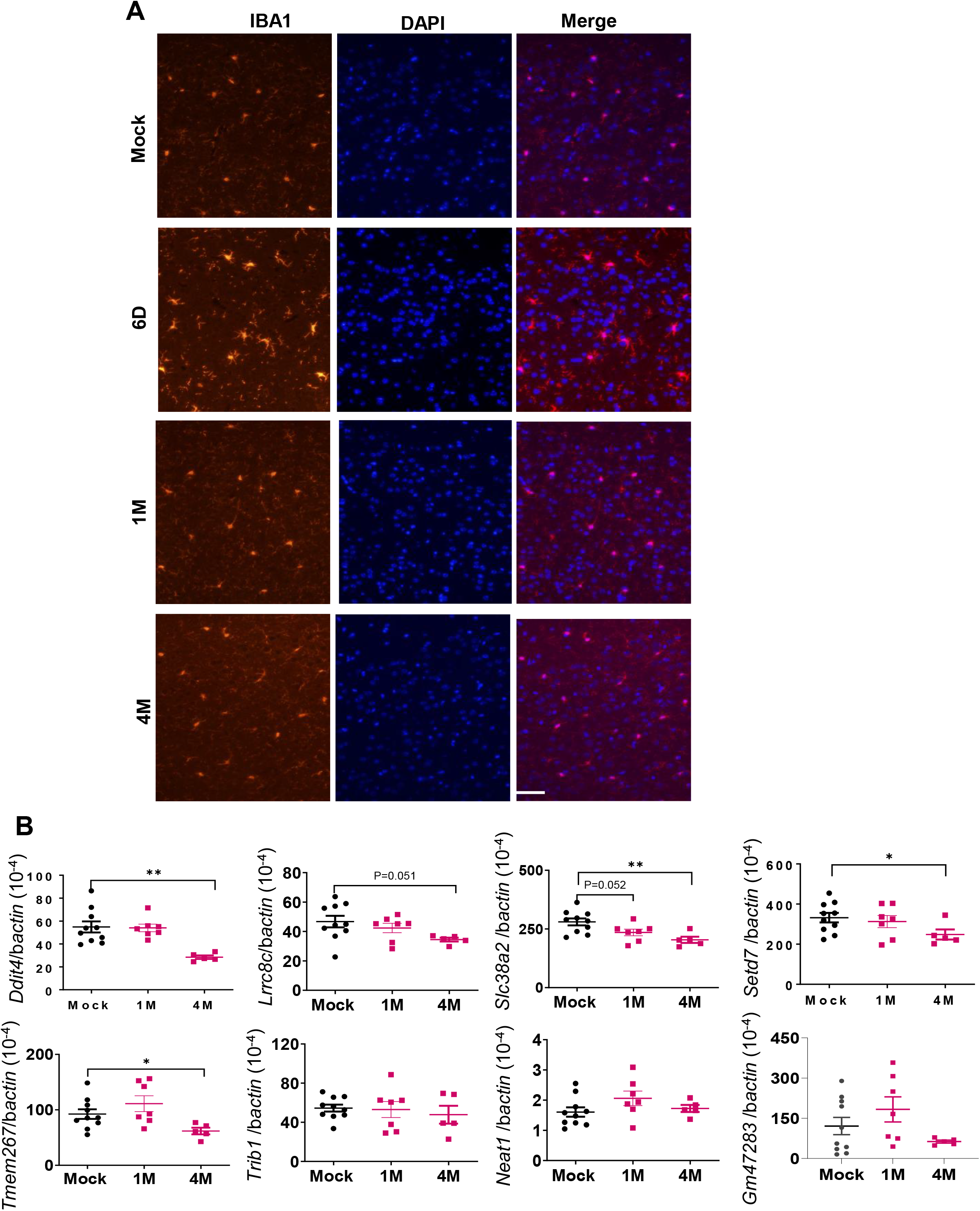
Microglia were activated during acute SARS-CoV-2 infection in the brain but were minimally activated post-acute infection. **A.** Immunofluorescence images of brain tissues stained with DAPI (blue) and anti-Iba1 (red) at both acute infection (day 6, D6) and post-acute infection (1-month (M) and 4M pi). **B.** Q-PCR analysis of the levels of 8 differentially expressed genes identified by transcriptomic analysis in the brain samples of mock-or SARS-CoV-2-infected samples at 1M and 4M. n= 5 to 10. \*\**P* < 0.01, or \**P* < 0.05, compared to the mock group.

Blood and spleen tissues were collected to assess peripheral SARS-CoV-2-specific cellular and humoral immune responses in the surviving mice. At 1 month pi, splenic T cells, including both CD4^+^ and CD8^+^ T cell subsets, produced robust T helper (Th)-1 prone immune responses upon *in vitro* re-stimulation with S peptide pools compared to the mock group (**Fig. 7A-C**). Sera of the surviving mice also maintained high titers of RBD-binding IgG and neutralizing antibodies against Delta variant during the 4-month post-acute infection. Notably, comparable levels of neutralization activity against Omicron strain were detected in the Delta-variant infected mice compared to those infected with Omicron strain at 4 months (**Fig. 7D-E**). Overall, these results suggest that mice surviving SARS-CoV-2 Delta variant infection developed long-lasting Th1 and antibody responses in the periphery post-acute infection.

**Figure 7.**
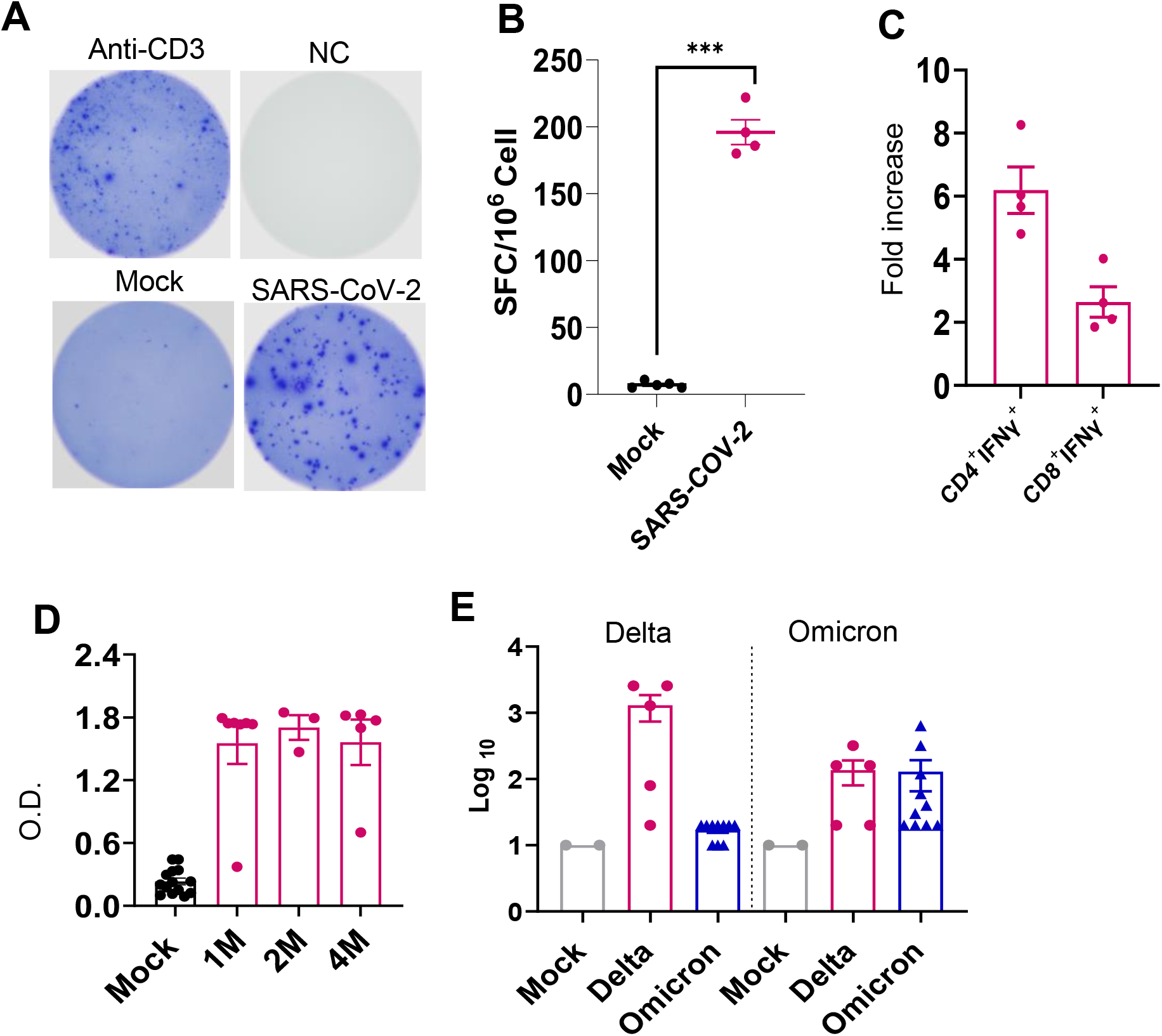
SARS-CoV-2 Delta variant induced persistent systemic cellular and humoral immune responses post-acute infection. K18-hACE2 mice were infected with a sublethal dose of SARS-CoV-2 Delta variant**. A-C.** At 1 month pi, splenocytes were collected from surviving mice and mock group to measure T cell responses. **A**-**B.** ELISPOT quantification of splenic T cell responses. Splenocytes were stimulated with SARS-CoV-2 S peptides, anti-CD3, or blank (negative control, NC) for 24 h. **A.** Images of wells of T cell culture. **B.** Spot forming cells (SFC) were measured by IFN-γ ELISPOT. Data are shown as # of SFC per 10^6^ splenocytes. n= 5. **C.** Splenocytes were cultured *ex vivo* with S peptide pools for 6 h, and stained for IFN-γ, CD3, CD4, or CD8. Fold increase of IFN-γ^+^ CD4^+^ and CD8^+^ T cells expansion compared to the mock group is shown. **D.** Sera SARS-CoV-2 RBD-binding IgG titers at 1 month (M), 2M and 4M pi. O.D. values were measured by ELISA. n = 5 and 10 for Delta-variant infected and mock respectively. **E.** At 4M pi, sera neutralizing activity against SARS-CoV-2 Delta variant or Omicron B.A.2 variant was measured by plaque reduction neutralization test (PRNT). mNG-NT_50_ titers are shown, n =2, 5, and 10 for mock, Delta-variant or Omicron-infected group respectively. \*\*\**P* < 0.001, compared to the mock group.

## Discussion

It is known that high risk of PASC is associated with people with prior COVID-19 infection. More evidence suggest that frequencies of PASC symptoms increased with SARS-CoV-2 variants, in particular, the pre-Omicron variant compared to the original prototype virus infection ^3,30–32^. Here, we infected K18 hACE2 mice with SARS-CoV-2 Delta variant to recapitulate PASC in COVID19 patients. We found that SARS-CoV-2 Delta variant replicated productively in lung and brain and triggered robust local inflammatory responses during the acute phase of infection in K18-hACE2 mice. Weight loss, neuroinflammation, and mortalities were observed during acute infection. Surviving mice showed no additional weight loss, viral clearance, and minimal neuroinflammation but persistent neuropsychiatric state and motor-associated behavior changes for months post-acute infection.

Our longitudinal behavior studies confirm ataxia and cognitive dysfunction in SARS-CoV-2 variant-infected mice post-acute infection. The behavior studies suggest that the neuropsychiatric state and motor behavior of surviving mice remain impaired or deteriorated for months pi, whereas reflex and sensory functions appear to recover over time. These findings align with a recent 2-year retrospective cohort study that reported an increased risk of psychotic disorder, cognitive deficit, dementia, and epilepsy or seizures persisted in long COVID patients ^41^. Furthermore, downregulation of *Ddit4*, *Slc38a*2, *Tmem267m*, *Lrrc8c*, and *setd7* expression levels in the brain at 4 months pi suggests ataxia, impairment of memory, synaptic plasticity, motor, and cognitive abilities, neuronal dysfunction and degeneration in the surviving mice^37–40^. RNA-seq analysis revealed activation of immune pathways in the CNS, including “complement activation pathway,” “phagocytosis recognition,” and “humoral immune response mediated by circulating immunoglobulin” at 1 month and 4 months pi. The transcriptome results support neurological behavior changes observed in the surviving mice. These findings also indicate that complement-dependent engulfment of synapses may lead to the cognitive dysfunction. The spike protein and its fragment has been reported to be able to cross the blood brain barrier and enter the CNS^42^, and is directly involved in COVID-19 induced cognitive dysfunction ^43^ via complement-dependent engulfment of synapses in mice ^44^. No detectable levels of S1 in the CNS were found during post-acute phase in this study. Although we noted neuroinflammation and microglia reactivity during acute infection, minimal to mild microglia activation was found months post-acute infection. It’s likely that viral infection and/or viral fragment entered in the CNS during the acute phase induced inflammation and caused neuronal injury which resulted in neurological sequelae. Interestingly, we noted that surviving mice maintained potent protective Th-1 prone and humoral immune responses in the periphery post-acute infection. The role of peripheral immune factors in SARS-CoV-2 variant-induced PASC pathogenesis will be investigated in future studies.

Several rodent models have been used to study long-COVID ^45–47^. In line with a recent report in a golden hamster model of long-COVID, we found detectable infectious virus in the CNS during the acute infection phase accompanied with behavioral changes at 1 month after viral clearance ^46^. Induction of CCL11 expression in the periphery tissues and CNS during acute COVID19 infection similar as reported in a mild-respiratory COVID model in immunocompetent mice via delivery of AAV vector to express human ACE2 to the trachea and lungs ^47^. Furthermore, results from the transcriptome analysis align with the impaired neurogenesis findings reported in both studies. Nevertheless, we did not note microglia reactivity, neuroinflammation, and induction of proinflammatory cytokines in the CNS post-acute phase as reported in both studies. Both prior studies are limited to 4 weeks to 7 weeks post-acute infection and the use of wild-type prototype virus. Another study reported increased reactive astrocytes and microglia, hyperphosphorylated TDP-43 and tau, and decrease in synaptic protein synaptophysin-1 and defective neuronal integrity in A/J mice 12 months post-infection. Although the study recapitulates long-term sequelae of COVID-19, a mouse hepatitis virus 1 (MHV-1) was used in this study^48,49^.

In summary, our results suggest that infection in K18-hACE2 mice recapitulates the persistent clinical symptoms reported in long COVID patients. Our immunological and transcriptomic analysis provides new insights into the pathogenesis of the disease. The K18-hACE model of long-COVID may be useful to evaluate efficacy for future development of novel SARS-CoV-2 vaccines or therapeutics.

## Supporting information

SinghAdam Supple Figures

## Supplementary Figure Legends

**Supplementary Fig. 1. Viral loads in brain and periphery tissues at acute and post-acute infection.** K18-hACE2 mice were infected with a sublethal dose of SARS-CoV-2 Delta variant, or PBS (mock). Viral loads in brain (**A**), lung (**B**), liver (**D**), kidney (**E**), and blood (**F**) tissues at indicated days (D) or months (M) pi were measured by Q-PCR. **C.** S1 protein levels in brain lysates were measured by ELISA. Data are presented as means ± SEM. \*\*\**P* < 0.001, \*\**P* < 0.01, or \**P* < 0.05 compared to mock.

**Supplementary Fig. 2. SARS-CoV-2 Delta variant infection induces minimal or mild inflammation in the brain and lung post-acute infection. A.** Histopathology of Delta variant-infected or mock-infected brain Cortex and brain stem at 1 month (M) or 4M pi. Views of representative areas from each group show no inflammation is observed in the control or infected brains. Bar = 80 μm. **B.** Cytokine and chemokines expression levels in the lungs at 1M and 4M were measured by Q-PCR. Data are presented as the fold increase compared to mock-infected mice (means ± SEM). n= 5 to 10. \*\**P* < 0.01, or \**P* < 0.05 compared to mock.

**Supplementary Fig. 3. RNA-seq analysis of SARS-CoV-2-infected mouse brain shows genes with upregulation of immune signaling and enrichment of immune signaling related pathways.** Representative GSEA plots showing enrichment of immune pathways based on RNAseq results of brain samples collected from 1-month (**A**) and 4-month (**B**).

## ACKNOWLEDGEMENTS

We thank Dr. Linsey Yeager for assisting in manuscript preparation. This work was supported in part by NIH grants R01AI127744 (T.W.), R01 NS125778 (T.W.), and R01AI176670 (T.W.), and a Pan Pilot Grant at UTMB (P.K.).

## AUTHOR CONTRIBUTIONS

A.S., A.A., J.Z., V.K. performed the experiments. A.S., A.A., I.C.; T.W. designed the experiments; A.S., A.A., B.P., X. Y., V.K., W.J., P.Y.S., and T.W. analyzed the data; P.K. and T.W. secured the funds for the project; T.W. wrote the initial draft of the manuscript, and other coauthors provided editorial comments.

