## Supplementary figures and images for "A Murine Model of Post-acute Neurological Sequelae Following SARS-CoV-2 Variant Infection"

### SinghAdam Supple Figures

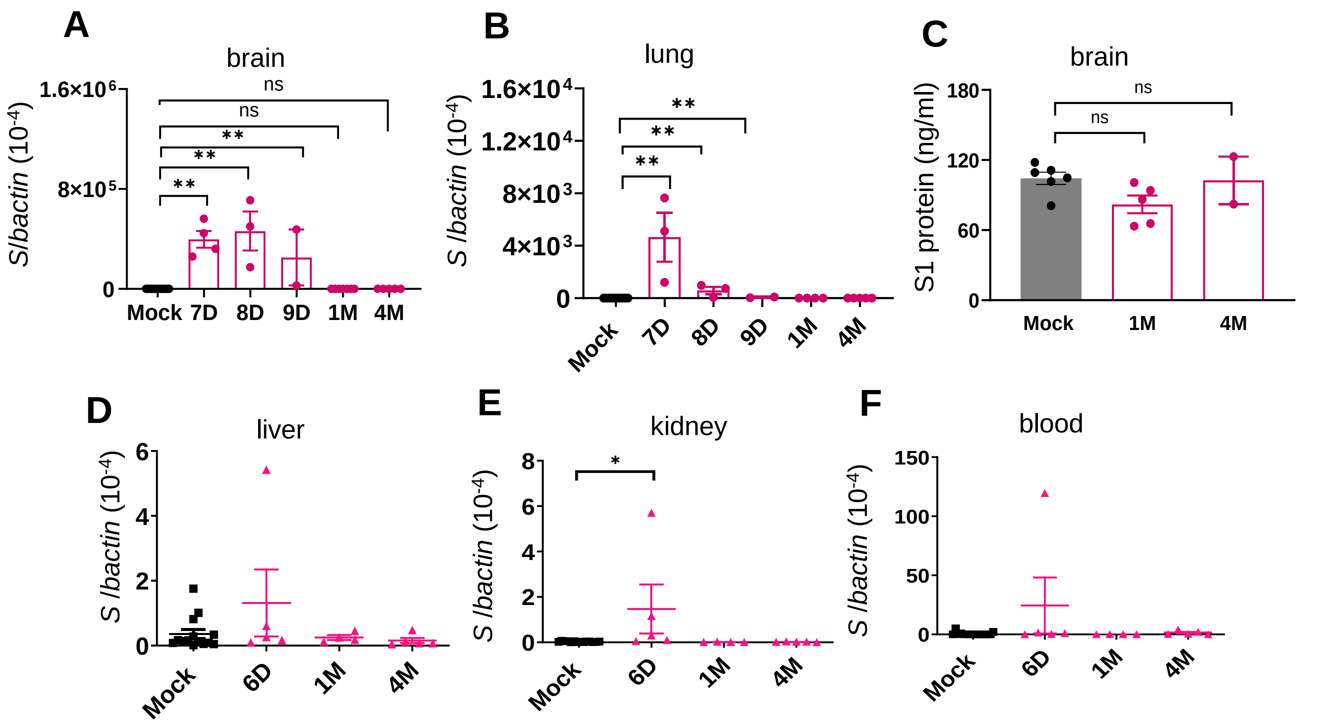


**Supplementary Figure 1**

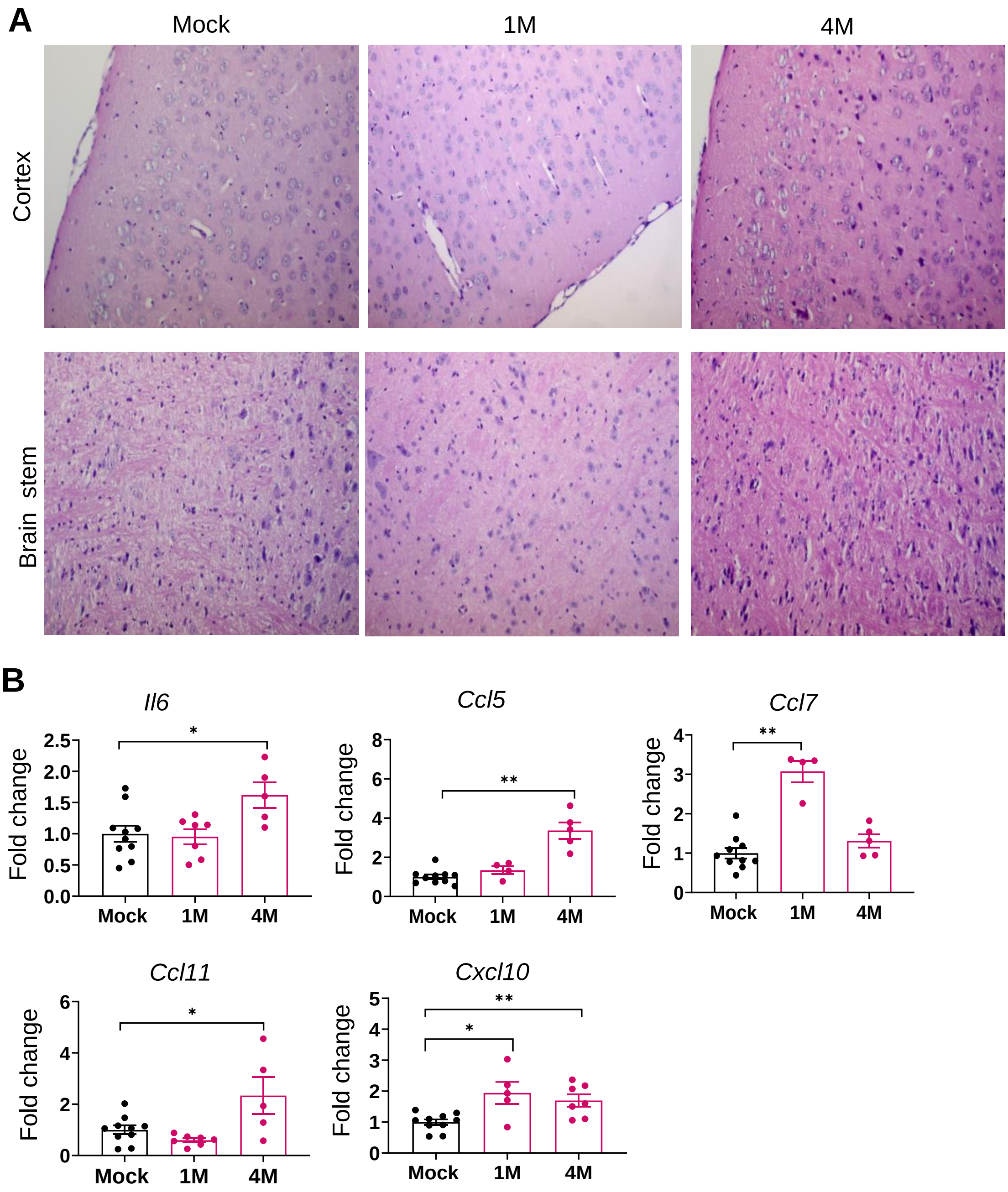


**Supplementary Figure 2**

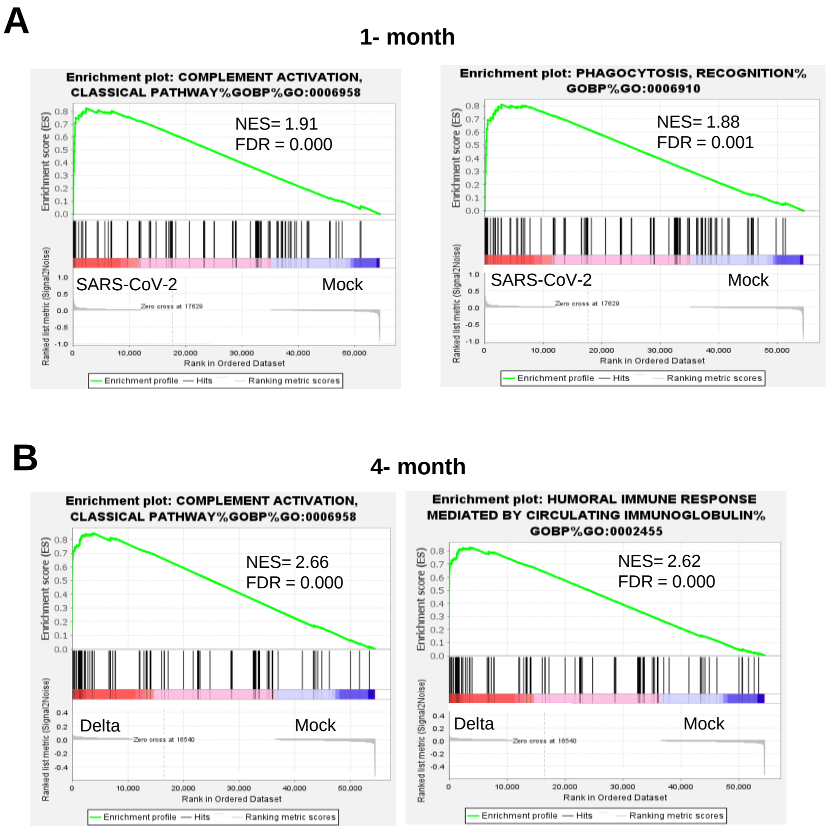


**Supplementary Figure 3**
