## Supplementary material for "A Murine Model of Post-acute Neurological Sequelae Following SARS-CoV-2 Variant Infection": Table S1

**Supplementary Table 1: Sequence of qPCR primers: for validation of RNAseq analysis**

| **Gene** | **Primer sequence** | **Reference** |
| --- | --- | --- |
| *Neat1* | Forward: GTTCCGTGCTTCCTCTTCTG  Reverse: GTGTCCTCCGACTTTACCAG | ^1^ |
| *Lrrc8c* | Forward: AACTCGGTCACCGGAATCAT  Reverse: CCCCAGAGATTAATGTGGCT | ^2^ |
| *Trib1* | Forward: GGAAGTTCGTCTTCTCCACCGA  Reverse: GCAGCCATGTTTATCTGACAGCG | <https://www.origene.com/catalog/gene-expression/qpcr-primer-pairs/mp217531/trib1-mouse-qpcr-primer-pair-nm_144549>  NCBI Reference Sequence:  NM_144549.4 |
| *Slc38a2* | Forward: TAATCTGAG CAATGCGATTGTGG  Reverse: AGATGGACGGAGTATAGCGAAAA | ^3^ |
| *Tmem267* | Forward: GGCAGTAGTCACTGGAATCAGG  Reverse: CTTCGCGGAAGAGTCAAAGCGG | <https://www.origene.com/catalog/gene-expression/qpcr-primer-pairs/mp211555/tmem267-mouse-qpcr-primer-pair-nm_001039244>  NCBI Reference Sequence: NM_001039244.4 |
| *Setd7* | Forward: TTGACGGAGAGATGCTCGAAGG  Reverse: GAAGGAGAGCATCGCTGGAGAT | <https://www.origene.com/catalog/gene-expression/qpcr-primer-pairs/mp215317/setd7-mouse-qpcr-primer-pair-nm_080793>  NCBI Reference Sequence: NM_080793.6 |
| *Ddit4* | Forward: ACTGCGAGTCCCTGGACAGCA  Reverse: TTGGCACACAGGTGCTCATCCT | <https://www.origene.com/catalog/gene-expression/qpcr-primer-pairs/mp203836/ddit4-mouse-qpcr-primer-pair-nm_029083>  NCBI Reference Sequence: NM_029083.2 |
| *Gm47283* | Forward: CTGAGAAGAGGGCGTCAGAT  Reverse: CTGTCAGAGTGAAGGGCAGA | ^4^ |
